# Integrative system biology analysis of barley transcriptome – hormonal signaling against biotic stress

**DOI:** 10.1101/2021.10.19.464927

**Authors:** Zahra Soltani, Ali Moghadam, Ahmad Tahmasebi, Ali Niazi

**Author notes:** **Author Contributions** Zahra Soltani analyzed the data and write the text, Ali Moghadam designed the topic and reviewed the text, Ahmad Tahmasebi reviewed the text, Ali Niazi reviewed the text.

## Abstract

Biotic stresses are environmental factors that cause a variety of crop diseases and damages. In contrast, crops trigger specific transduction signaling pathways that the hormones are the central players. Integrative OMICS for systems genetic engineering approach contributes in the understanding of molecular mechanisms. In this research, the system biology approaches were applied to discover particular molecular interactions between biotic stresses and hormonal signaling in barley. The meta-analysis of the data identified a total of 1232 and 304 differentially expressed genes (DEGs) respectively so that were significantly involved in defense processes and hormone signaling. A total of 24 TFs belonged to 15 conserved families and 6 TFs belonged to 6 conserved families were identified for biotic and hormonal data respectively, whereas NF-YC, GNAT, and whirly families were the most abundant groups. The functional analysis of the upstream regions for over-represented cis-acting elements revealed that were involved activation of transcription factors in response to pathogens and hormones. Based on the co-expression analysis, 6 and 7 distinct co-expression modules related to biotic stresses and hormonal signaling were respectively uncovered. The gene network analysis also identified novel hub genes such as *TIM10, DRT101, ADG1*, and *TRA2* which may be involved in regulating defense responses to biotic stresses. In addition, many new genes with unknown function were obtained. Since this study represents a first preliminary curated system biology analysis of barley transcriptomic responses to biotic stresses and hormone treatments, introduces important candidate genes that may be beneficial to crop biotechnologists to accelerate genetic engineering programs.

## Introduction

Cereals are directly related to the food source and energy supply and are constantly faced with a wide range of biotic stresses that negatively effect on their survival and growth (Sharma *et al*., 2013). These stresses are caused by living organisms such as bacteria, viruses, fungi, nematodes, and pests (B. Liu et al., 2014; Rastgoo & Alemzadeh, 2011). Among biotic stresses fungal pathogens are involved in 70 to 80% of plant diseases (Arabi *et al*., 2021; Ray *et al*., 2017). The cereal barley (*Hordeum vulgar*) is very susceptible to fungal diseases (Massey *et al*., 2007) like *Fusarium graminearum* that is a serious trouble in wet or semiwet climates (McMullen *et al*., 1997; Parry *et al*., 1995). These pathogens are a major limitation to global crop production; hence, plant genes encoding resistance to pathogens, are important tools for combating disease (Chakraborty & Chakraborty, 2021; Milne *et al*., 2019; J. Wang *et al*., 2021). Resistance conferred by these genes is preferable over using commercial fungicides (Singh *et al*., 2015). In addition to these pathogens, one of the most important pest on cereals is the bird cherry-oat aphid (*Rhopalosiphum padi* L.), which reduces plant growth without specific leaf symptoms (Luo *et al*., 2019). Breeding of barley for *R. padi* resistance has been revealed that there are several resistance genes, decrease aphid growth. Several gain-of-function and loss-of-function mutants have showed genes related with fundamental defense and effector-triggered immunity (Jones & Dangl, 2006; Shen *et al*., 2007; D. Wang *et al*., 2006).

Access to OMICS information on the behavior of defense genes against pathogens or pests helps to decipher the gene networks that led to detection of pathogen specific responses (Reymond *et al*., 2000). Large-scale data generated from functional genomic studies on the plant-pathogen interactions have been provided massive information about the expression patterns of key genes and interactions (Kaul *et al*., 2000). In recent years, OMICS data and progressive statistical analyses such as computational systems biology have collected exquisite opportunities to dominate biological complexity (Bolhassani *et al*., 2021; Tahmasebi *et al*., 2019b).

Meta-analysis and network analysis are two potent strategies that integrate OMICS data to recognize core gene sets that regulate the complex traits (Frierson Jr *et al*., 2002; Sharifi *et al*., 2018). These approaches provide a robust statistical framework for re-evaluating key findings, improving sensitivity by increasing sample size, testing new hypotheses, (Z. Liu *et al*., 2013; Rung & Brazma, 2013) and characterizing of novel gene candidates in plant breeding programs (Ashrafi-Dehkordi *et al*., 2018; Shaar-Moshe et al., 2015; Tseng *et al*., 2012). Therefore, progress in whole-genome transcriptome analysis techniques have analyzed gene regulation and plant-pathogen interactions in various conditions, allowing identification of stress-specific biomolecular networks and hormonal signaling pathways. For example, the main response to pathogens may be due to the cross-talks among hormone-related pathways (Pieterse *et al*., 2009) such as ethylene (ET), jasmonic acid (JA), salicylic acid (SA) and abscisic acid (ABA), and/or signaling due to calcium (Dodd *et al*., 2010; Schulz *et al*., 2013), activated oxygen species (ROS) (Torres *et al*., 2006), and phosphorylation chains (Asai *et al*., 2002). Transcription factors transmit these signaling indications and activate or suppress the expression of genes involved in immune responses and metabolic processes. Thus, most plant immune responses are transcriptionally controlled (Tao *et al*., 2003; Wise *et al*., 2007).

In the other words, plants are equipped with an array of defense strategies to protect themselves against pathogens, one of which is the ability to prime defense (Dey *et al*., 2014). Depending on the location of the initial infection and the virulence of the attacker, this type of induced resistance is often referred to as induced systemic resistance (ISR) or systemic acquired resistance (SAR). ISR is caused on the colonization of plant roots by nonpathogenic soil microbes and protects aerial tissues of dicots and monocots from necrotrophic pathogens and pests (Balmer *et al*., 2013; Pieterse *et al*., 2012; Walters *et al*., 2013). On the other hand, SAR is induced in systemic and uninfected tissues of a plant on prior foliar pathogen challenge and is mainly beneficial against biotrophic pathogens (Fu & Dong, 2013; Vlot *et al*., 2009). Many hormonal cross-talks are involved in both defense systems for example, induction of SAR is more through SA while ISR requires JA and ET signaling pathways (Van Loon *et al*., 1998). These accumulating signaling molecules moderate the defense responses and when used exogenously, are enough to induce the resistance (Ryals *et al*., 1996). The protection mediated by ISR is mainly less than that obtained by SAR (Van Loon, 2000) and a degree of dependence on plant genotype is observed in the production of ISR (Bloemberg & Lugtenberg, 2001). However, ISR and SAR together present a better protection than each of them alone, and this shows that they can act increasingly in inducing resistance to pathogens (Van Wees *et al*., 2000).

In this study, in order to identifying the hub genes and gene networks and providing further insight into the mechanisms involved in the response of the barley to various pathogens and hormonal signal transductions, meta-analysis and co-expression network analysis were applied on barley transcriptome data in response to pathogens and hormones. These genes were identified through functional enrichment analysis of metabolite pathways, transcription factors (TFs), protein kinases (PKs), and microRNA families. A more accurate knowledge of barley-pathogen interactions might be helpful for plant biotechnologists to produce the pathogen-tolerant cultivars.

## Methods

### Data collection

In this study, the raw microarray expression data of barley exposed to diverse biotic stresses including fungi, bacteria, viruses, insects, and hormones including JA, gibberellic acid (GA), methyl jasmonic acid (mJA) were extracted from publically available databases of Gene Expression Omnibus (http://www.ncbi.nlm.nih.gov/geo) and Array Express (http://www.ebi.ac.uk/arrayexpress). The data selection was done based on the “Minimal Information about a Microarray Experiment” (MIAME) requirements. Therefore, the studies with relatively similar genetic background without any mutated or transgenic samples, with suitable quality and biological and technical replicates of controls and treatments were chosen (Cohen & Leach, 2019). In addition, the six most critical elements contributing towards MIAME including raw data, normalized data, sample annotations, experimental design, array annotations, and data protocols were considered (Brazma *et al*., 2001). The total of 10 studies including 479 biotic stress samples (**Table S1**) and 4 studies including 46 hormonal treatment samples (**Table S1**) were selected for the data analysis. The raw data of all the studies were downloaded in two platforms, Affymetrix (Accession: GPL1340) and Agilent (Accessions: GPL14904 and GPL15513). The probe-gene maps and probe-annotation files were downloaded from the Affymetrix site (http://www.Affymetrix.com).

### Pre-processing and normalization

Low quality data is responsible for the differences between tests, which may be due to lack of standards as well as inadequate testing methods, statistical analysis, validation, and/or reporting of the studies (Dupuy & Simon, 2007; Jafari & Azuaje, 2006; Shi *et al*., 2006). Therefore, the normalization of the Affymetrix microarray raw expression data for each array was performed with robust multichip average (RMA) algorithm (Irizarry *et al*., 2003) using the Expression Console software (V. 1.3.1). After background correction, RMA performs a quantile normalization were carried out and then uses median polish as summarization method (Durinck, 2008). Agilent microarray data (two colors) were processed using Flex Array software (V. 1.6.3). At the first, we performed the background correction using normexp algorithm and MLE method to estimate the intensity of expression of raw microarray data (Ritchie *et al*., 2007; Silver *et al*., 2009). Finally, the normalization between arrays was performed using the quantile method (Bolstad, 2001) to reduce the variance and probe set differences between arrays. In addition, Agilent microarray data (one color) were processed in the R program using the package LIMMA and LOESS method (Smyth, 2005).

### Meta-analysis and identification of DEGs

In this study, the controls and treatments for each study were separately defined, and guardianship data in the form of Uniprot IDs as input files for meta-analysis were integrated using R package metaDE. The merged datasets were filtered again with the exception of 20% of the expressionless genes (with low expression intensity) and 20% of the non-educational genes (genes with quantitative changes). Finally, the results based on Rank Prod method, which is a non-parametric method were used. In this method, genes were ranked according to their up- and down-regulation, fold change (FC), and FDR. In this way, the Rank Prod algorithm calculates pairwise FC with replicates for each gene between treatment and control samples in both directions, respectively, and converts FC into rank among all genes, then searches for genes that are continuously top ranked across replicates (Hong & Breitling, 2008). Therefore, up- and down-regulated genes obtained with FC cutoff of log2 FC >1 or log2 FC <−1 and *FDR* < 0.05 were only considered for the meta-analysis (Balan *et al*., 2017).

### Functional annotation and enrichment analyses

The DEGs selected by the meta-analysis were further analyzed and characterized. Gene ontology (GO) enrichment analysis of the DEGs to find important pathways was conducted using the Kyoto Encyclopedia of Genes and Genomes (KEGG) pathway and DAVID (http://david.abcc.ncifcrf.gov/) platforms (Du *et al*., 2010). GO enrichment was performed on the hypergeometric statistics followed by a FDR < 0.05 correction for multiple comparisons mapped to each category (Shaar-Moshe et al., 2015). GO study was performed on the unique list of TAIR IDs of the genes obtained in non-redundant stress data. The data sets were log transformed and significant genes were selected according to *P-value* < 0.05. Where data for multiple time points were available, the first time point representing the maximum genomic response was selected. Values obtained from databases were an averaged value for each gene that represented the overall transcript change across time points.

### Identification of TFs, PKs and miRNAs families

The iTAK database (http://itak.feilab.net/cgi-bin/itak/index.cgi) was used to identify the transcription factors (TF) and protein kinase (PK) families of the barley DEGs. In the next step, the psRNATarget database (http://plantgrn.noble.org/psRNATarget) was used to identify possible miRNAs among DEGs. Then, to further demonstrate the response to biotic stresses and hormones, all of stress-responsive miRNAs were collected and compared with the enriched GO identifier to annotate all genes. WEGO software (http://wego.genomics.org.cn/) was used for plotting GO annotation results

### Identification of modules

In order to network analysis, R package WGCNA (Weighted Gene Co-expression Network Analysis) was conducted on the matrix of normalized expression values of DEGs for all samples to identify groups of DEGs with similar expression patterns. For this purpose, the meta-analysis output file was merged with the initial normalized data and then the treatment and control samples were separated in the columns. Thus, WGCNA file was prepared based on the meta-analysis expression values. A resemblance matrix was calculated based on Pearson correlations between each DEG pair and converted into an adjacency matrix by applying a power function. Then followed by, the topological overlap matrix (TOM) (Yip & Horvath, 2007) was calculated for hierarchical clustering analysis. Finally, a dynamic tree cut algorithm was used to identify gene co-expression modules. This method can detect nested clusters and have been shown to better detect outliers (Langfelder *et al*., 2008). Finally, the modules were defined with a minimum module size of 30 genes and a threshold of 0.3 and a minimum module size of 100 genes and a threshold of 0.99 for biotic and hormonal data respectively.

### Identification of hub genes

Network visualization and calculation of topological properties for each module were performed using the Cytoscape software (V. 3.6.1). Hub genes were also screened using plug-in of Cytoscape, CytoHubba (Chin *et al*., 2014). To identifying the hub genes (30 nodes with the highest interaction) in the network, computational algorithms of Maximal Clique Centrality (MCC) were used as the most effective method (Li *et al*., 2020). Then all hub genes were studied for enrichment analysis in terms of Molecular Function (MF), Biological Process (BP), and Cellular Component (CC) by DAVID (http://david.abcc.ncifcrf.gov/) database.

### Protein-protein interactions

Annotation and analysis of individual data were accomplished using Network Analyzer (Xia *et al*., 2014), a web-based tool for protein-protein interaction network analysis and visual exploration. The list of biotic stress and hormonal related genes was determined. The unique list of homologous TAIR IDs of each data groups was BLASTed against the STRING interactome database with default parameters (minimum required interaction score = 0.150). The low confidence was selected to make simpler network and to study the important and key connectivity’s. Finally, the network was drawn by Cytoscape software.

### Cis-elements analysis

In order to obtain the genomic sequences, 1000 bp from the upstream flanking region of DEGs, the genes were used to the BLASTX search against the Ensemble Plants (http://plants.ensembl.org). Then, the MEME (meme.nbcr.net/meme/intro.html) (Bailey et al., 2009) database was used to identify the conserved cis-acting elements. The motifs were counted and the abundance of regulatory elements in the promoter region of various genes was investigated. In the next step, we used Tomtom v 5.0.1 tool (http://meme-suite.org/tools/tomtom) (Gupta et al., 2007) to obliterate redundancy motifs and determine known CRE based on the motif database of JASPAR CORE 2018 plants (Khan et al., 2017) with threshold *E-value* cut-off of 0.05. GMO tool (http://meme-suite.org/tools/gomo) was also used to identify the biological roles of the cis-acting elements (Buske et al., 2010).

## Results

In this study, in order to identify key genes in response to pathogens and interaction with various hormones, we analyzed some microarray data of barley by system biology approaches. Meta-analysis approach integrated DEGs from microarray datasets which were expressed consistently with statistical significance. A schematic workflow, summarizing major steps of this study is shown in **Fig. 1**. In addition, co-expression network analysis was performed to identify correlated DEGs, so that genes with unique functions were integrated in common modules.

**Fig. 1.**
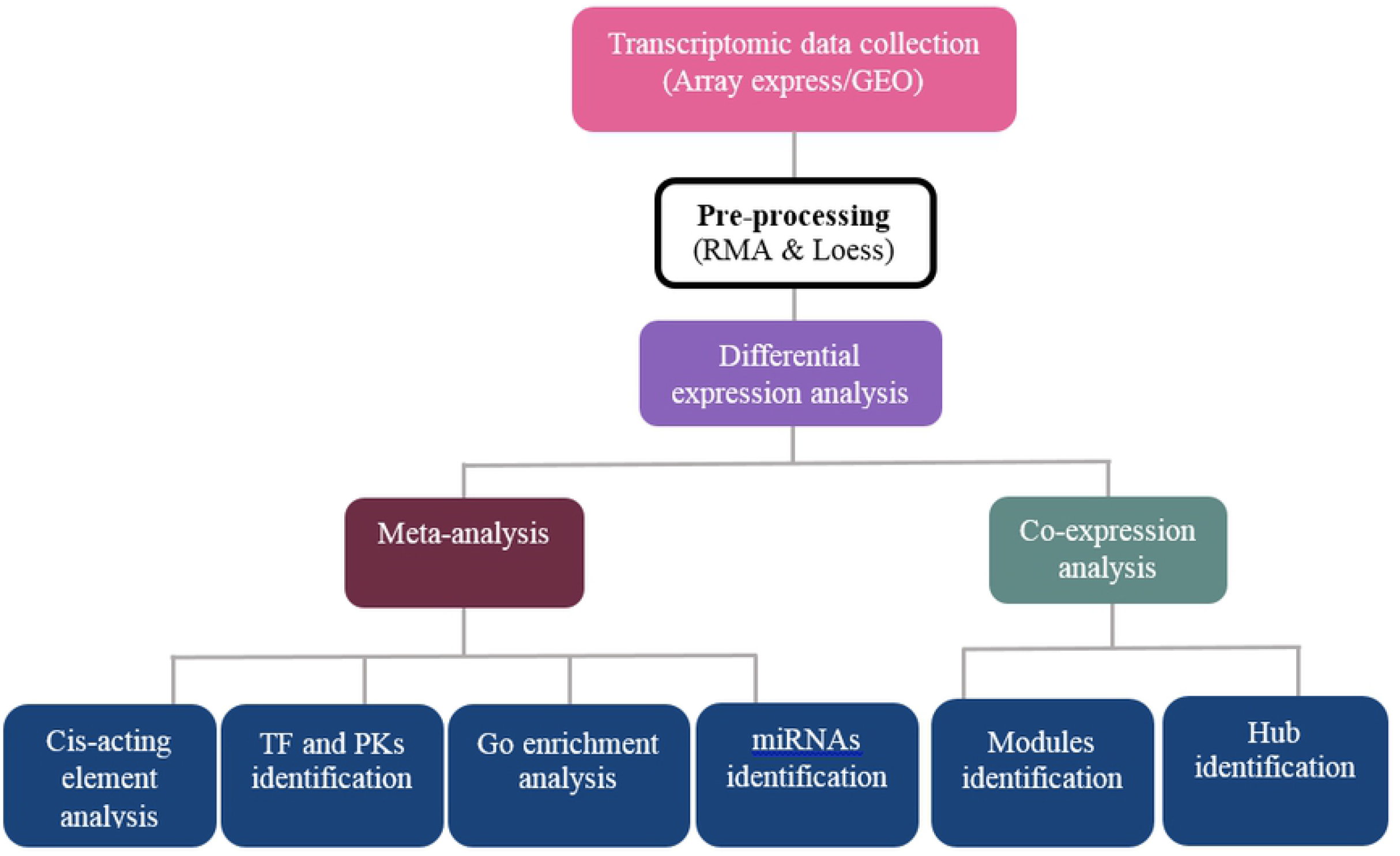
Schematic overview of the integrative strategy for understanding aspects of response of barley to biotic stresses and hormonal treatments.

### Pre-processing and normalization

The purpose of pre-processing microarray gene expression data is to select a small set of genes which can be used to improve the attention and performance of classification from a high dimensional gene expression dataset (Hu *et al*., 2006). Pre-processing is used to detect data noise and reduce the impact of existing noise on the device learning algorithm. Raw microarray data pre-processing involves image processing, quality assessment, background correction, normalization, summarization, and then high-level analysis such as gene selection, classification, or clustering (Freudenberg, 2005). Reducing heterogeneity among studies is a necessary step for direct combination of expression data in differential expression meta-analysis. To reduce the any systematic differences for meta-analysis, an integrative approach of within-study RMA and LOESS normalization were developed and then the samples were aggregated to remove duplicates. Fig. S1 compares the heterogeneity among samples of different studies after different normalization steps. The proposed framework of normalization and standardization reduced the batch effects and noises, and facilitated direct merging of samples from different experiments (**Fig. S1**).

### Meta-analysis and identification of DEGs

In this study, Rank Prod method was used to analyze microarray data and identify DEGs for biotic stress and hormonal treatment studies (**Table S1**). Although the Fisher’s combined probability test adds the transformed logarithm *P-values* obtained from individual studies, under the null hypothesis, and follows a chi-squared distribution with 2 k degrees of freedom, where k is the number of studies being combined (Rhodes *et al*., 2002),the Rank Prod, calculates the product of the rank of pair-wise differences between each biological sample in one group versus another group during the studies (Breitling & Herzyk, 2005) and calculates the product of the ranks of FC in each inter-group pair of samples (Hong *et al*., 2006). Therefore, in the Rank Prod method could only be accomplishment with up- and down-regulation analysis separately. Because theoretically, it is easy to modify the algorithm to analyze up- and down-regulation simultaneously (Tseng et al., 2012).

Based on the results, the meta-analysis was able to find 1232 genes, which were significant differentially expressed (adjusted *P-value* and FDR < 0.05). Among the biotic data, 546 up-regulated genes and 642 down-regulated genes were identified, and 177 up- and 127 down-regulated genes were identified for hormonal data (**Table S2**). Finally, we created a list of up- and down-regulated genes and compared to identify common and specific genes. Venn diagrams showed the numbers of specific and common regulated genes between the two types of biotic stresses and hormonal treatment studies (**Fig. 2-3**).

**Fig. 2.**
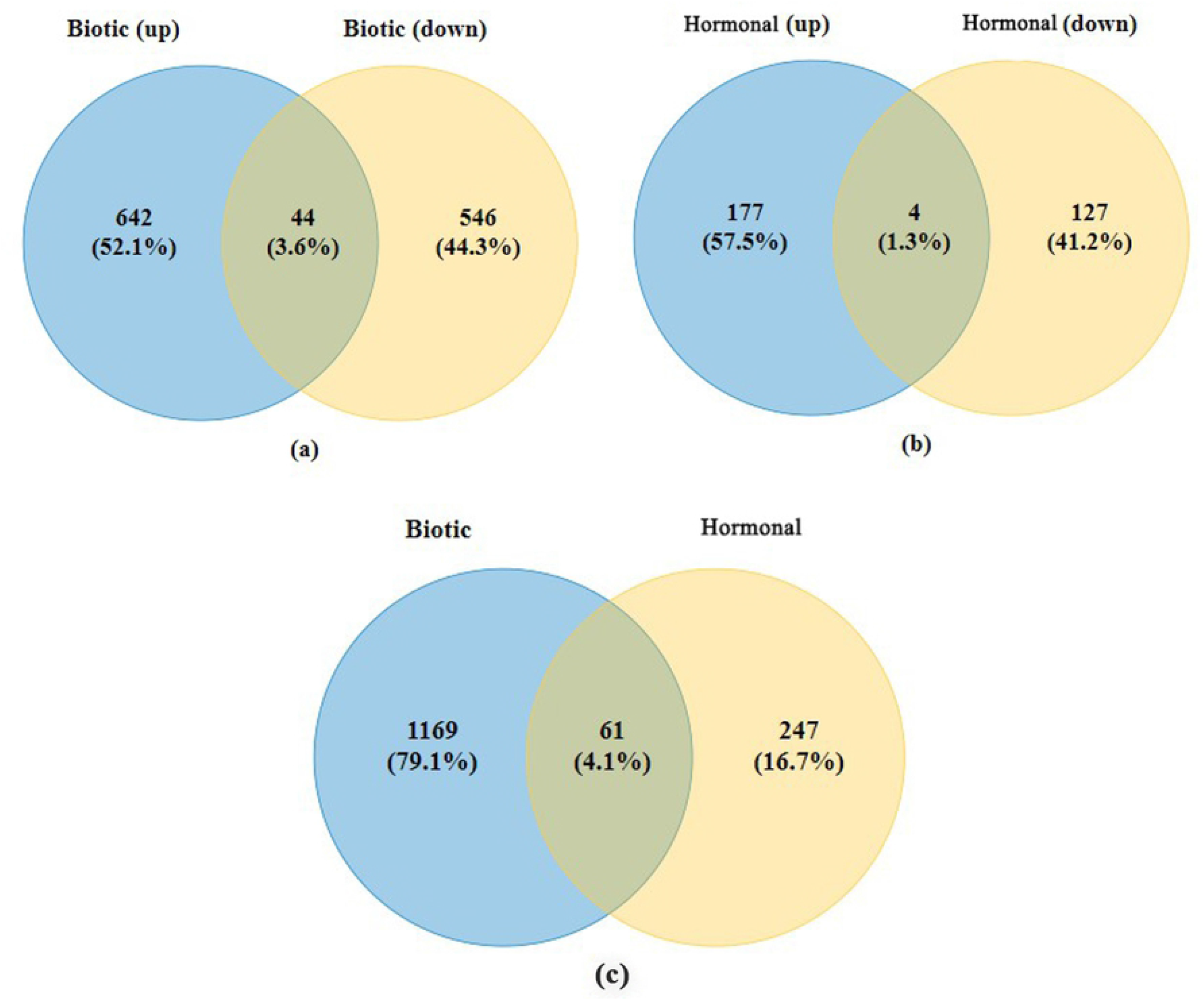
Venn diagrams of DEGs inferred from biotic and hormonal studies. a) Up- and down-regulated biotic DEGs. b) Up- and down-regulated hormonal DEGs. c) Total DEGs of biotic and hormonal studies.

**Fig. 3.**
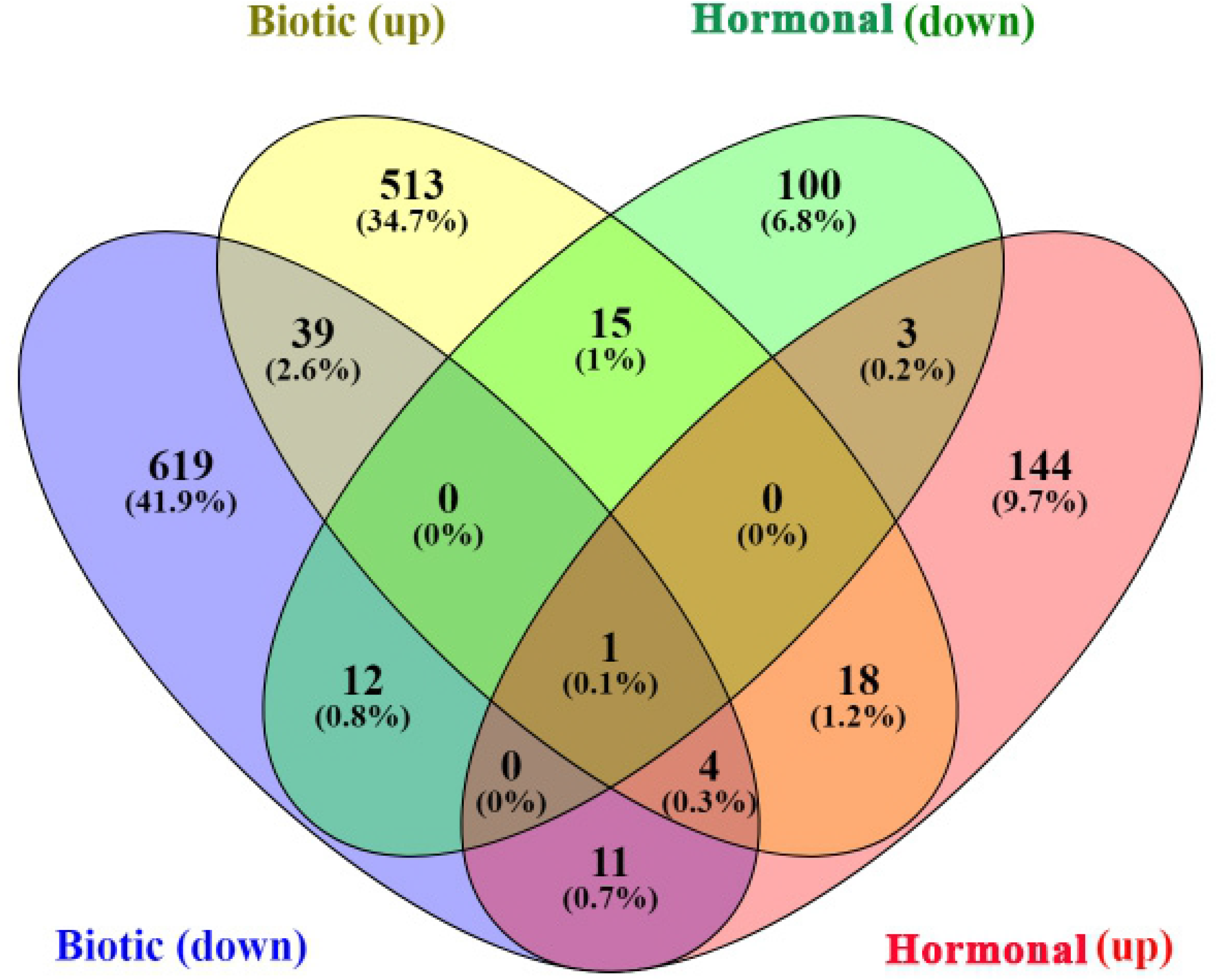
Venn diagrams of DEGs inferred from biotic and hormonal studies. d) Comparison of up- and down-regulated DEGs in both biotic hormonal studies.

A total of 61 common DEGs between biotic and hormonal data were identified in which 18 DEGs such as *EMB1144, PAL1, GAD, ALDH7B4, JAZ1, NINJA* and some genes withunknown function including *AT1G22410, AT1G06550*, AT5G58950, and AT1G68300 were identified. In addition, a total of 12 DEGs that were significantly down-regulated including *MT3, SIR, ABCA2, GAMMA-VPE*, *APE1* as well as some genes with unknown function including *AT1G22410*, *AT3G15810, AT2G21180* and *AT4G09890* were identified.

### Enrichment analysis and functional annotation of DEGs

The 1232 and 308 DEGs related to biotic and hormonal studies respectively were subjected to GO and gene network analysis to explore other possible functions. The results of the GO in the DAVID database were divided into three different categories, BP, CC, and MF. The results of the biotic stress studies showed the highest percentage of DEGs involved in BP (**Fig. 4a**).

**Fig. 4.**
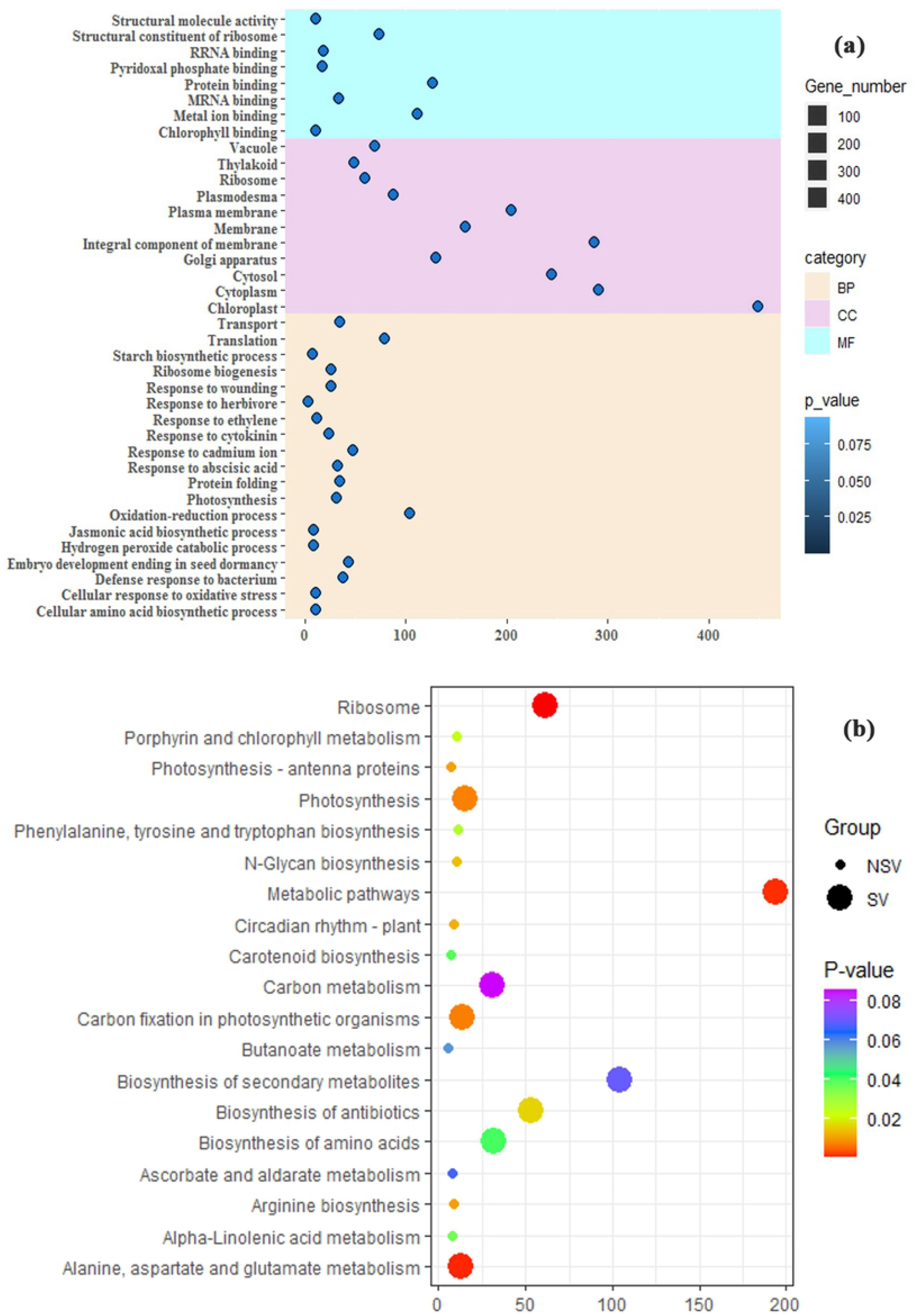
(a) Gene-ontology analysis of biotic DEGs. GO enrichment analysis of DEGs was retrieved using DAVID and the results were plotted using R software. The most significantly (P < 0.05) enriched GO terms in Biological Process (BP), Molecular Function (MF) and Cellular Component (CC) branches related to defense response are presented. The horizontal axis shows the number of genes based on the p-value, while the vertical axis represents Go-term include categories of MF, CC, and BP, respectively. (b) KEGG pathway analysis of biotic DEGs. Y-axis shows the pathways associated with biotic stress studies and the X-axis shows the amount of DEGs enriched in the pathway. The colors of each bubble are determined based on the *P-value* of the identified pathways. Y-axis label represents pathway, and X-axis label represents rich factor (rich factor = amount of differentially expressed genes enriched in the pathway/amount of all genes in background gene set). Size (group) and color (based on p-value) of the bubble represent amount of differentially expressed genes enriched in the pathway and enrichment significance, respectively. The Fig. was drawn using R language.

These DEGs were related to processes of oxidation-reduction process (8.4% genes), translation (6.4% genes), response to cadmium ion (3.9% genes), embryo development ending in seed dormancy (3.5% genes), protein folding (2.8% genes), and system of transport (2.8% genes). The most important mechanisms were related to response to ABA (2.6% genes), photosynthesis (2.5% genes), defense response to bacterium (2.4% genes), cytokinin (1.9% genes) and ET (0.9% genes), cellular response to oxidative stress (0.8% genes), JA biosynthetic process (0.7% genes), hydrogen peroxide catabolic process (0.7% genes), response to reactive oxygen species (0.4% genes). In the category of CC the top enriched GO terms included chloroplast, cytoplasm, integral component of membrane, and cytosol. In the category of MF, chlorophyll and protein binding, metal ion binding, structural constituent of ribosome were dominant. The KEGG pathway analysis for DEGs revealed a significant enrichment in pathways associated with biosynthesis of secondary metabolites, ribosome, metabolic pathways, and biosynthesis of antibiotics and amino acids (**Fig. 4b**). In addition, for hormonal studies in the most percentage of DEGs (**Fig. 5a**) involved in BP. The pathways of oxidation-reduction process (15% genes), defense response (5.5% genes) and responses to oxidative stress (3.2% genes) were the top enriched GO terms. In the category of MF, metal ion binding and oxidoreductase activity were dominant. In addition, the category of CC and KEGG pathway analysis were the same as the components of biotic DEGs (**Fig. 5b**).

**Fig. 5.**
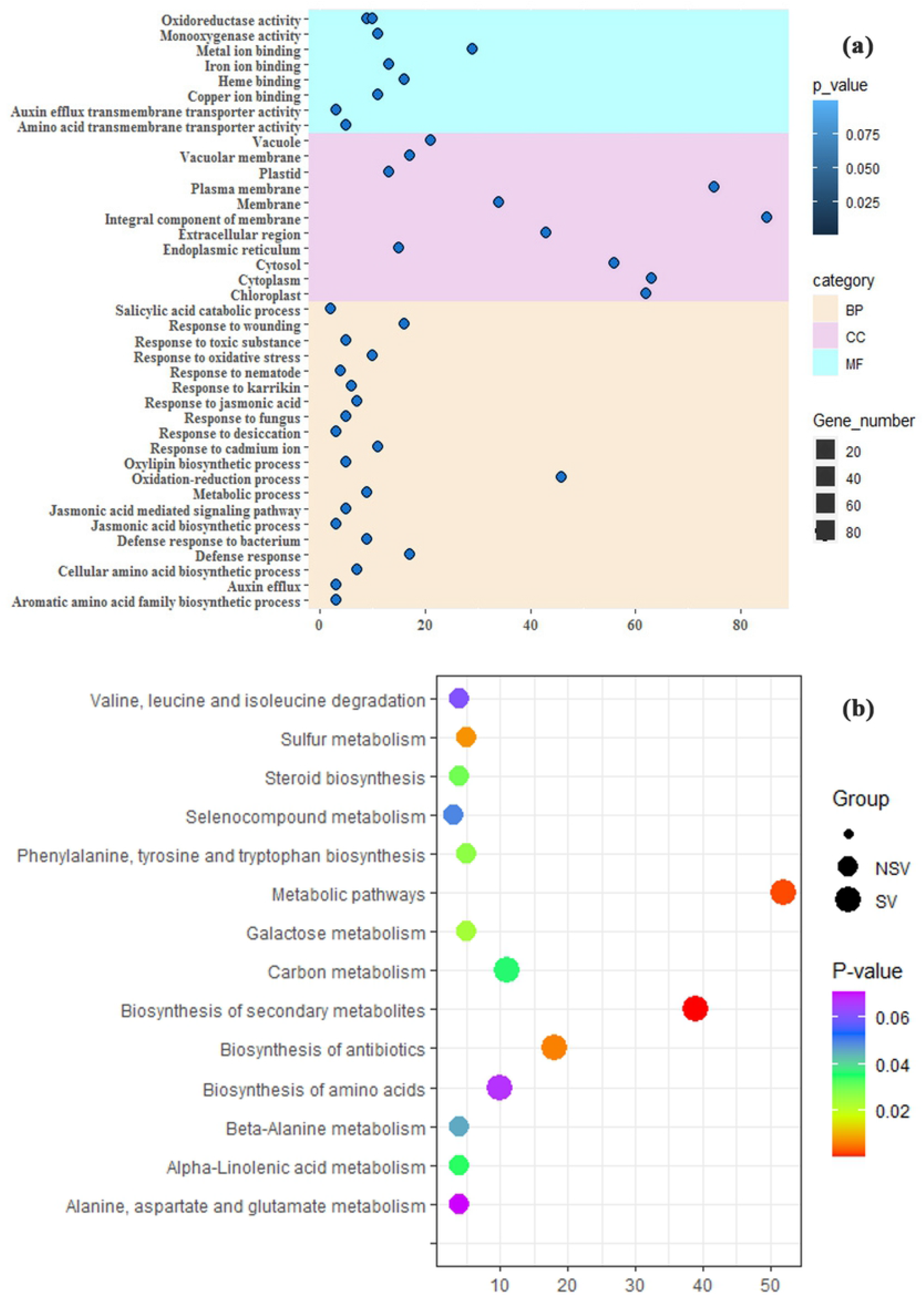
(a) Gene-ontology analysis of hormonal DEGs. GO enrichment analysis of DEGs was retrieved using DAVID and the results were plotted using R software. The most significantly (*P-value* < 0.05) enriched GO terms in biological process (BP), molecular function (MF) and cellular component (CC) branches related to signaling pathways are presented. The horizontal axis shows the number of genes based on the *P-value*, while the vertical axis represents Go-term include categories of MF, CC, and BP, respectively. (b) KEGG pathway analysis of hormonal DEGs. Y-axis shows pathways associated with hormonal stress and the X-axis shows the amount of DEGs enriched in the pathway. The colors of each bubble are determined based on the *P-value* of the identified pathway. Y-axis label represents pathway, and X-axis label represents rich factor (rich factor = amount of differentially expressed genes enriched in the pathway/amount of all genes in background gene set). Size (group) and color (based on *P-value*) of the bubble represent amount of differentially expressed genes enriched in the pathway and enrichment significance, respectively. The Fig. was drawn using R language.

### Co-expression analysis and module identification

In order to identify the different co-expressed modules, the R package WGCNA was applied on the 1232 biotic DEGs and 308 hormonal DEGs. Firstly, the network topology analysis was performed and appropriate power value was determined because the power value directly affected the average degree of connectivity and the independence within co-expression modules. When the power value was 10, the scale independence was up to 0.3 and 0.99 for biotic and hormonal data respectively that these were selected to produce a hierarchical clustering tree. Finally, the DEGs based on the dynamic tree cutting algorithm were grouped into 6 modules for biotic data ranged in size from 81 to 431 genes per module (**Fig. S2a, Table 1**) and 7 modules for hormonal data ranged in size from 141 to 926 genes per module (**Fig. S2b, Table 2**). In this way, WGCNA assigned a unique color label to each module that was used as specific module identifier in the analyses.

**Table 1.** Modules of biotic stress studies.

| Module name | Number genes | Module | Number genes |
| --- | --- | --- | --- |
| Blue | 296 | Blue.csv | 3858 |
| Brown | 229 | Brown.csv | 732 |
| Green | 81 | Green.csv | 42 |
| Red | 68 | Red.csv | 78 |
| Turquoise | 431 | Turquoise.csv | 9746 |
| Yellow | 124 | Yellow.csv | 2705 |

**Table 2.** Modules of hormonal treatment studies.

| Module name | Number genes | Module | Number genes |
| --- | --- | --- | --- |
| Blue | 926 | Blue.csv | 9181 |
| Brown | 305 | Brown.csv | 557 |
| Green | 206 | Green.csv | 505 |
| Red | 142 | Red.csv | 21 |
| Turquoise | 947 | Turquoise.csv | 8303 |
| Yellow | 221 | Yellow.csv | 394 |
| Black | 141 | Black.csv | 82 |

In the following, the hierarchical clustering of the TOM of DEGs related to both groups of the DEGs was illustrated (**Fig. S3**). As a result, an undirected weighted network with scale-free topology composed of modules of barley DEGs with correlated expression during pathogen infections was obtained. Finally, to understanding the biological functions associated with modules, the enrichment analysis for BP category was conducted (**Table S3**). The most significantly BP in Turquoise module were enriched in oxidation-reduction process and photosynthesis. The Blue module was associated with intracellular protein transport, Brown module with response to cadmium ion, Yellow module with translation, Green module with protein ubiquitination and Red module with response to stress. GO results related to hormonal modules (**Table S3**) Showed that the most important BP term enriched in Turquoise, Blue and Brown modules was oxidation-reduction process. In addition, a total of 600 common genes (17.1%) were identified between biotic stress and hormonal modules, which Turquoise (195 genes), Blue (169 genes), and Brown (117 genes) involved the most common genes.

### Hub genes

The network of the co-expressed modules was constructed to identify the genes with high connectivity or hub genes within each module. Because in biological network, each node represents a gene, which is linked to a different number of genes and is more important for its interaction with a large number of genes. A total of top 30 hub genes were chosen for each module. In the final, significant enrichment was performed for 177 hub genes from biotic modules and 198 hub genes from hormonal modules, and we were able to found functions that respond to biotic stress and hormones. The hub genes related to biotic modules were strongly enriched in photosynthesis, translation, and ribosome biogenesis pathways (**Fig. 6a**).

**Fig. 6.**
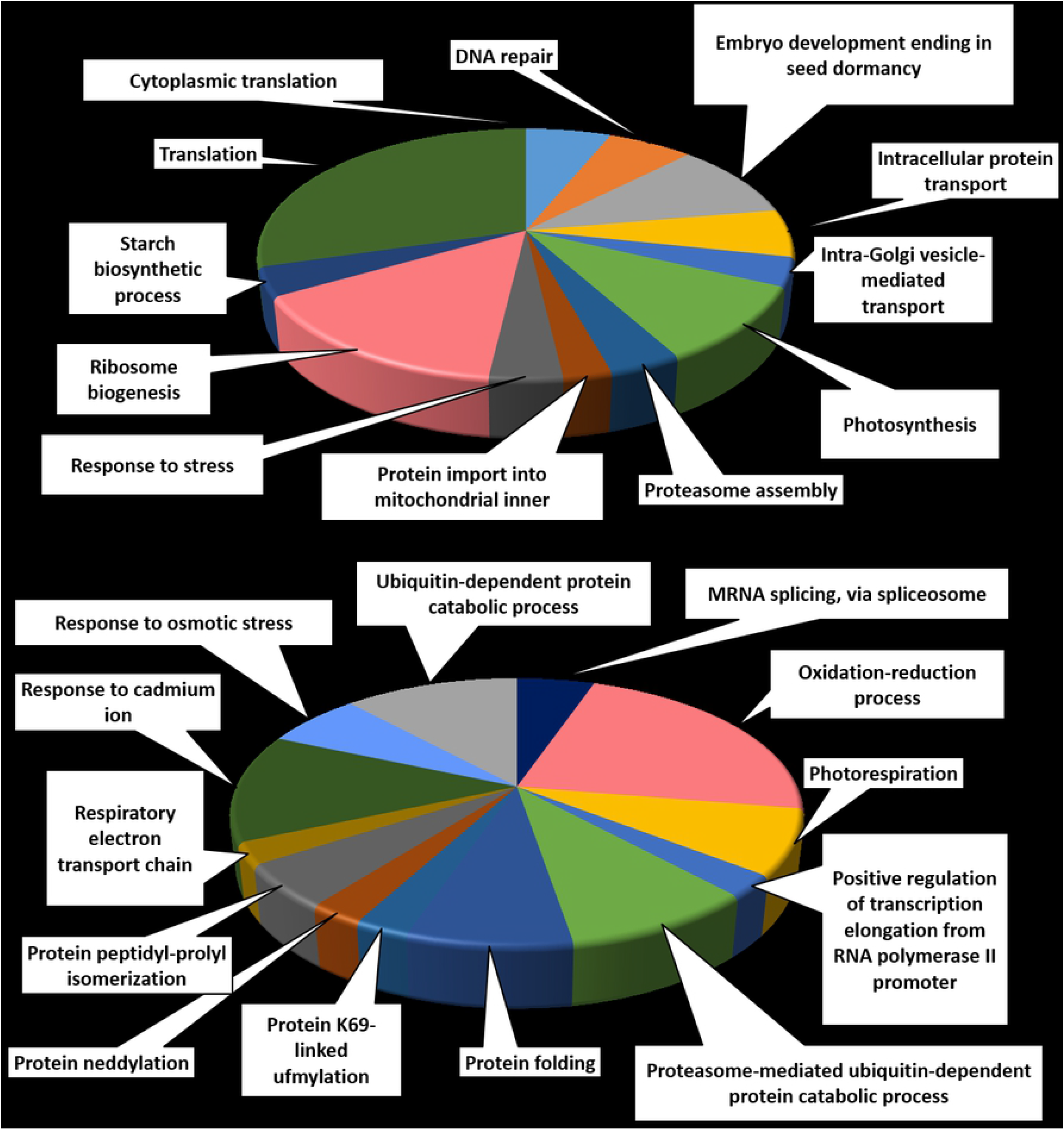
Distribution of GO enrichment of Biological Processes (BP) for hub genes in the two type of biotic stresses and Hormonal studies: (a) Biotic hub genes. (b) Hormonal hub genes.

In addition, among the hormonal modules, the top enriched BP terms were associated with oxidation-reduction process, response to cadmium ion, and ubiquitin-dependent protein catabolic process (**Fig. 6b**). In addition, a total of 10 common hub genes (2.7%) such as *NF-YC2*, SSI2, *ARAC1* and *PFD1* were identified between biotic stress and hormonal data (**Fig. S4**). In addition, some hub genes with unknown function including *AT3G26670, AT1G67620, AT1G05720, AT1G52600, AT5G59140* and *AT3G22290* were identified that may be considered potential candidates to be studied further.

### Identification of transcription factors

Because of the importance of transcription factors (TFs) as a powerful tool for the manipulation of complex network pathways, these genes were noticed. TFs are key players in stresses through transcriptional regulation. To identify transcription factors, DEG nucleotide sequences were used to the BLASTX search against the iTAK database. A total of 24 TF genes for biotic DEGs were found, which were belonged to 15 diverse TF families and directly or indirectly were involved in hormonal signaling and response to biotic and abiotic stresses (**Table S5, Fig. 7A**).

**Fig. 7.**
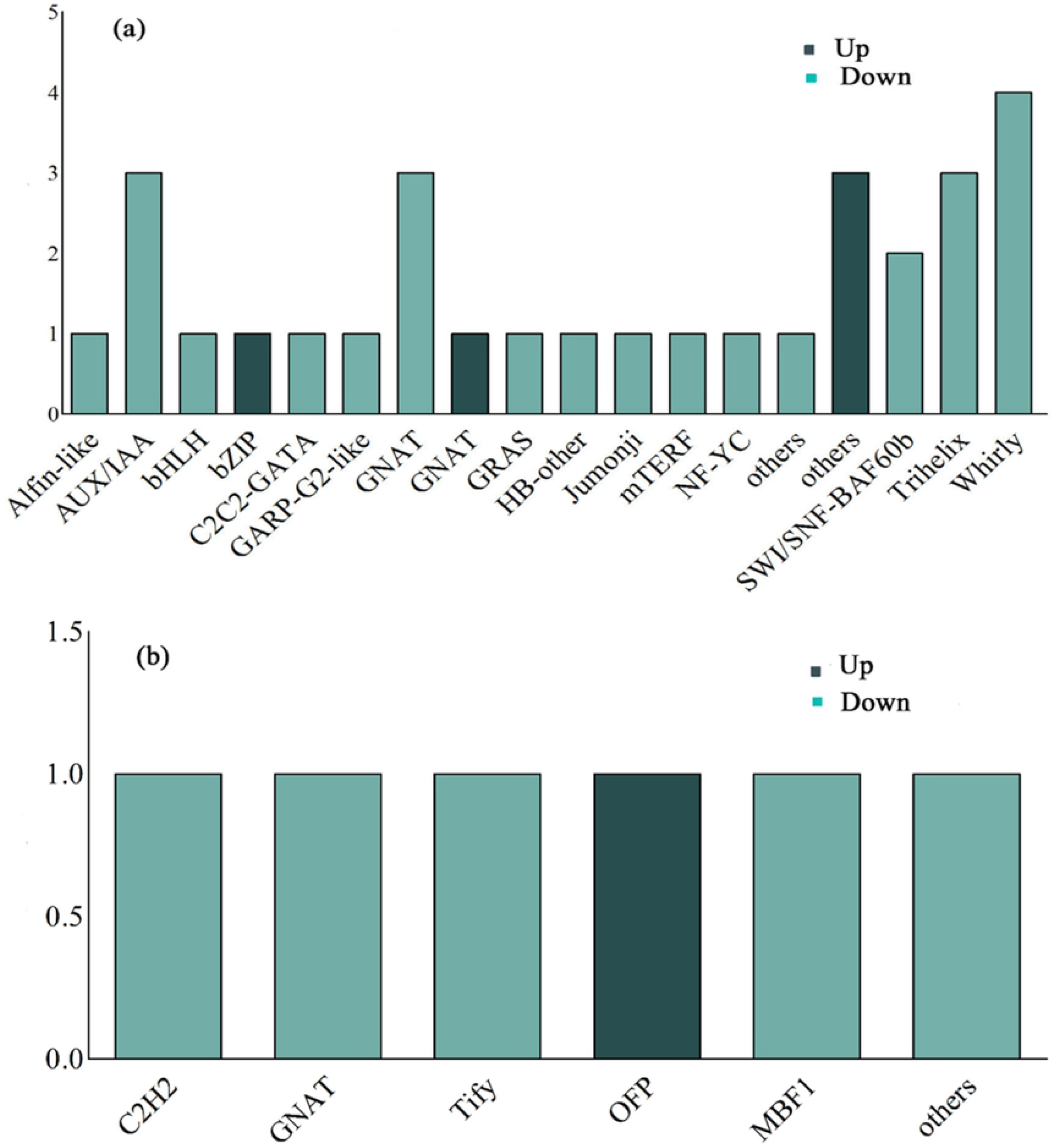
Distribution of TF families identified in DEGs. (a) The number of up- and down-regulated biotic TF families. (b) The number of up- and down-regulated hormonal TF families. The y axis refers to the number of genes. The x axis shown TF families.

Members of the AUX/IAA, GNAT, NF-YC and whirly families were the top classes. Among these transcription factor families, only 5 families were significantly up-regulated including bZIP and GNAT, while the most family members were down-regulated. These transcription factor families were more involved in oxidation-reduction process, response to wounding, defense response to bacterium, response to karrikin, response to ABA, JA mediated signaling pathways, and arginine biosynthetic process. In addition, among these transcription factors the Whirly (Turquoise module), GNAT, and NF-YC families (Green module) were also the hub genes (**Table S5**). In addition, a total of 6 TF genes related to hormonal DEGs belonged to 6 diverse TF families were identified (**Table S5, Fig. 7B**).

Among these transcription factor families only one family of GNAT was significantly up-regulated, while other families were down-regulated. These transcription factor families were more involved in response to oxidative stress, response to wounding, defense response to bacterium, response to ABA, JA mediated signaling pathways and defense responses. Brown module was enriched for Tify and OFP TF families whereas the Turquoise, Red and Black modules contained at least one member of GNAT and MBF1 families. Among the identified TF families of biotic stress and hormonal treatment studies, there was only one common gene. The significantly up-regulated *TIFY10A* (AT1G19180) was involved in response to JA, defense responses to bacterium, and regulation of defense responses.

### Identification of PKs

PKs are important signaling regulators in response to environmental stresses (Ho, 2015). To identify PKs, DEG nucleotide sequences were used to the BLASTX search against the iTAK database. Three PKs were identified among the biotic DEGs classified into RLK (Receptor-Like Kinase)-Pelle and TKL families (**Table 3**) that were totally up-regulated.

**Table 3.**
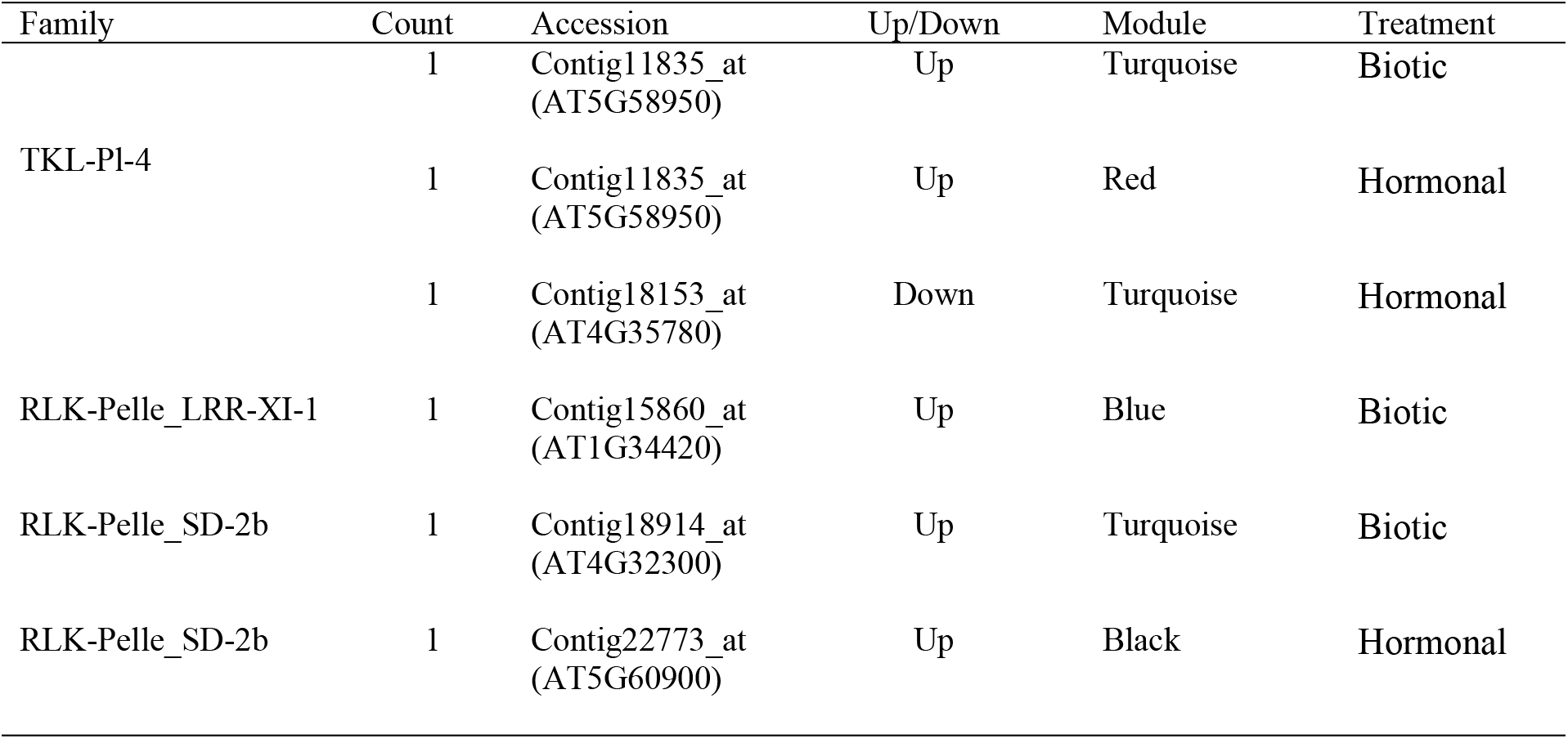
PKs for biotic stress and hormonal treatment studies.

The RLK-Pelle was the largest group among these families which was commonly identified in Turquoise and Blue modules. In addition, TKL-Pl-4 were identified in Turquoise module. Among hormonal DEGs, three PKS were identified, which were classified into RLK-Pelle and TKL families (**Table 3**). The RLK-Pelle were also identified in Black module only one of the TKLs is down-regulated and the rest of the PKs are up-regulated. Among identified PKs of biotic and hormonal DEGs, there was only one common gene. The significantly up-regulated *TKL-Pl-4* (*AT5G58950*) was the largest group among these families which was belonged to Red and Turquoise modules.

### Identification of miRNAs

To predict potential miRNAs that target the DEGs, psRNATarget tool was used. A total of 432 miRNAs belonged to 56 conserved families were found among the biotic DEGs (**Table S6**). In addition, a total of 85 miRNAs belonged to 32 conserved families were found among hormonal DEGs (**Table S6**). Among the identified miRNAs of both datasets, there were a total of 15 common target genes, which the miR6192 family comprised the highest frequency (**Fig. 8a**).

**Fig. 8.**
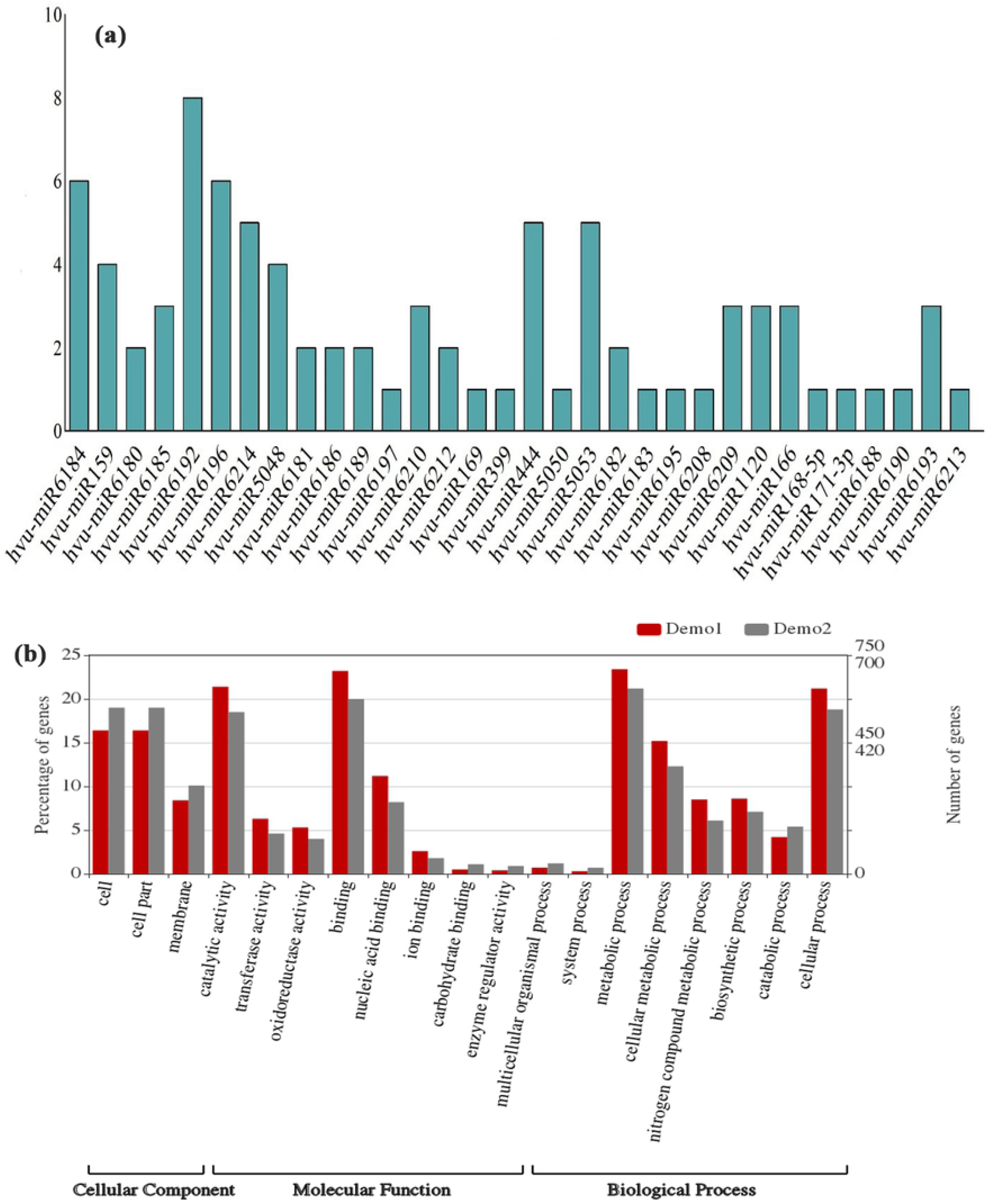
(a) Common miRNAs between biotic stress and hormonal studies. The y axis refers to the number of miRNAs. The x axis shown miRNA families. (b) Gene ontology analysis of miRNAs target genes. Categorization of miRNAs target genes with WEGO (http://wego.genomics.org.cn/cgi-bin/wego/index.pl) in biotic stresses and hormonal treatment studies. Notes: the horizontal axis is the GO classification; the ordinate is the percentage of gene number (left) and number of genes (right). This Fig. shows the gene enrichment of the secondary functions of GO in the background of the target gene and all genes of the differential expression miRNAs, reflecting the status of each secondary function in the two contexts. According to WEGO analysis, miRNAs can generally be divided into three categories: the cellular component, molecular function and biological process. All target: Demo1, De miRNA target: Demo2

All the putative target genes were categorized through GO analysis. The results showed that these genes were involved in different MFs, i.e., binding, oxidoreductase, transferase, and enzyme regulator activities, and played roles in many BPs, i.e., metabolic, cellular, and biosynthetic processes (**Fig. 8b**).

### Cis-acting elements

The 1000 bp upstream flanking region of DEGs was used to predict conserved consensus cis-regulatory elements (CREs). By MEME analysis, 11 significant motifs were identified with lengths ranging from 11 to 50 aa (**Tables 4 and 5**).

**Table 4.**
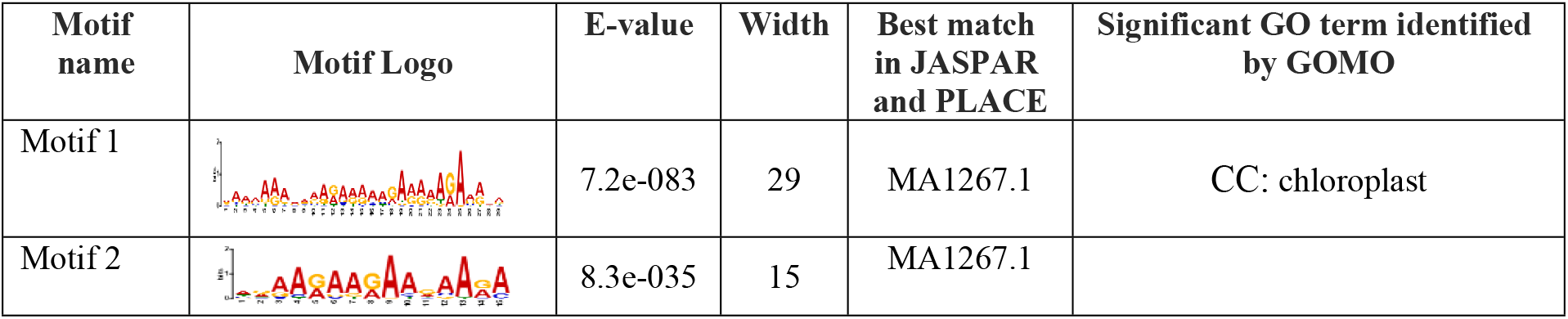
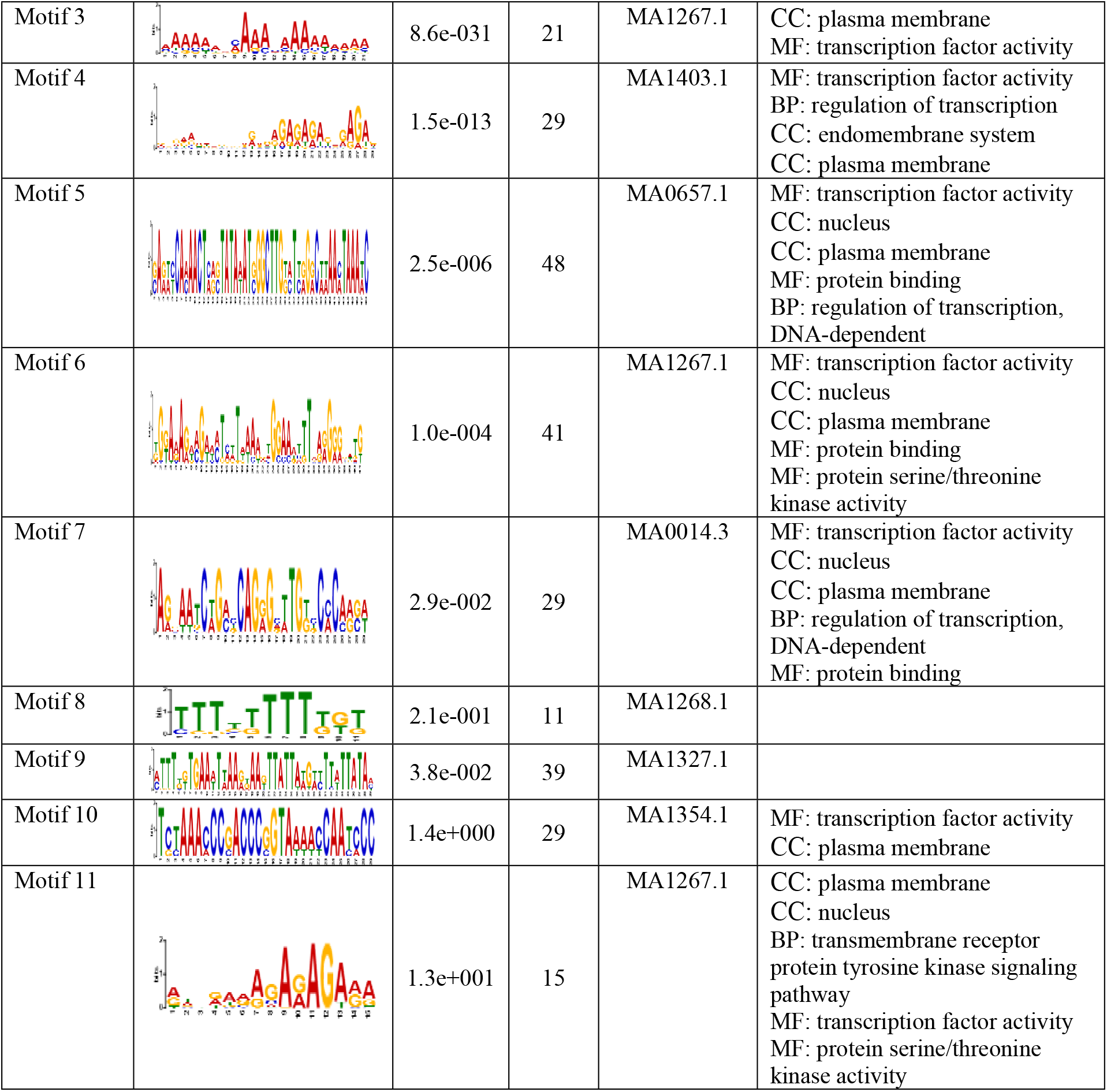
The conserved cis-acting elements found in promoter of biotic DEGs by the MEME analysis.

**Table. 5.**
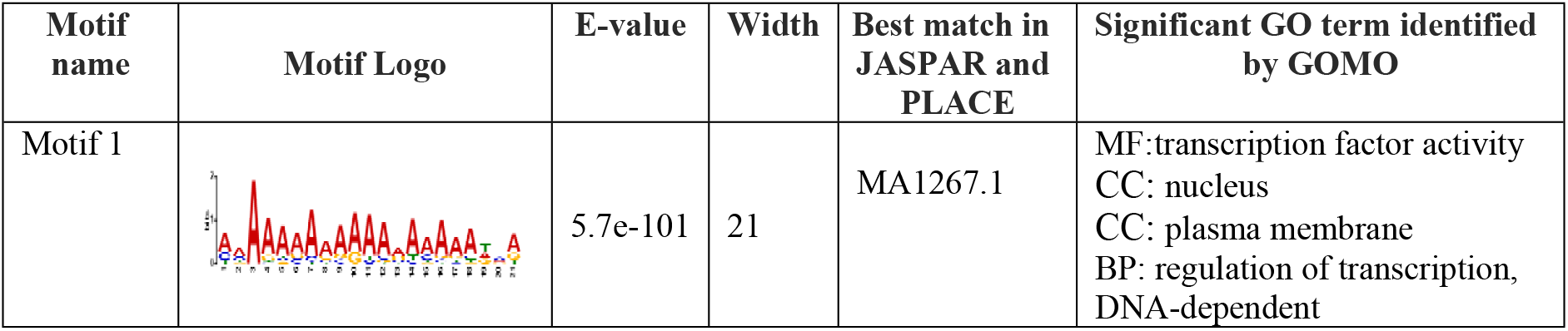
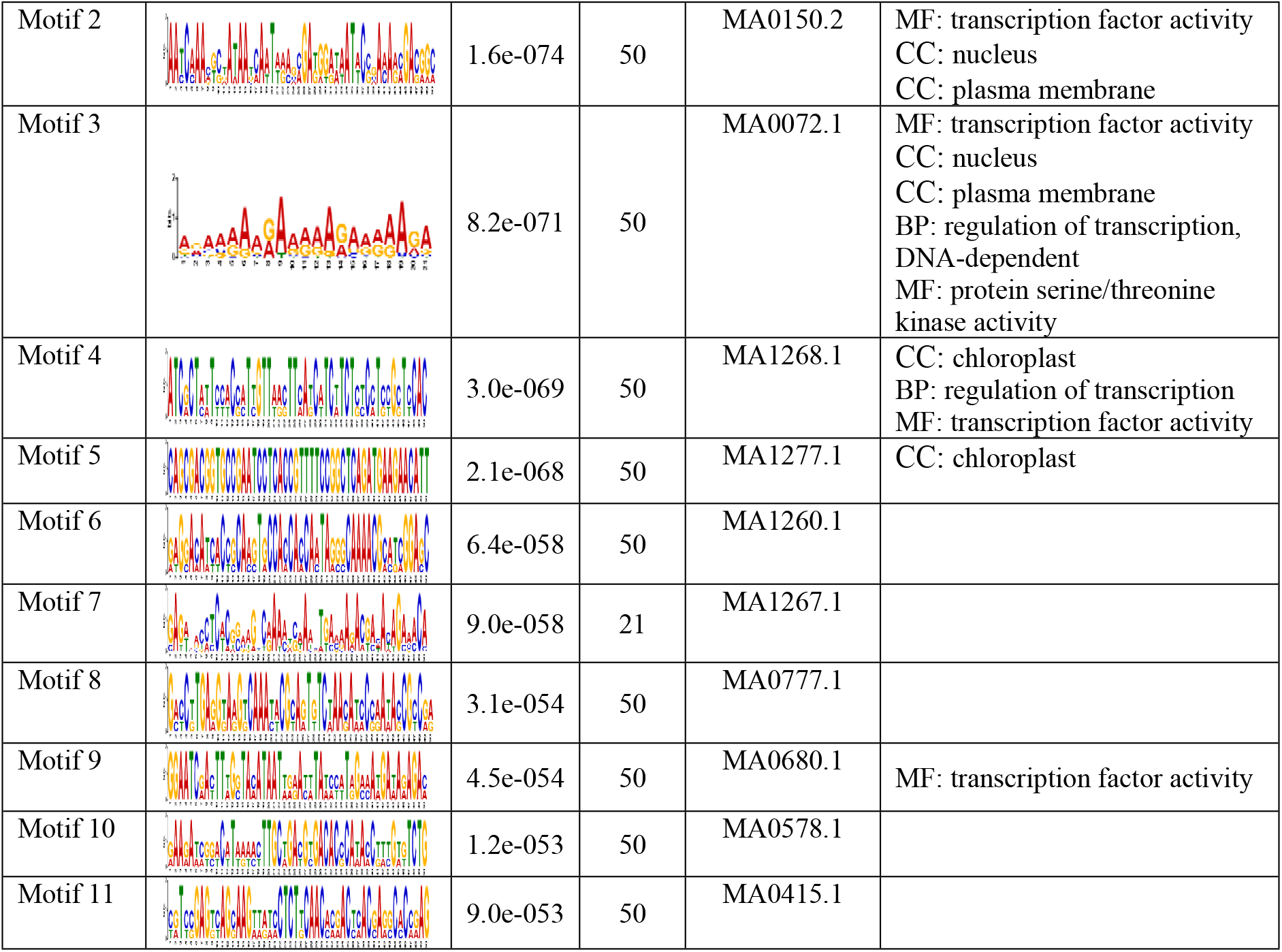
The conserved cis-acting elements found in promoter of hormonal DEGs by the MEME analysis.

After filtering (*P-value* < 0.05) motifs of 1 and 4 for biotic DEGs and motifs of 1 and 7 for hormonal DEGs showed the highest frequency. Moreover, the most of motifs are known as CREs which have related to C2H2 zinc finger factors (**Table S7**). The GOMO analysis for the motifs found by MEME detected many interesting BFs (**Tables 4 and 5, and Table S8**). GO on biotic DEGs indicated that these motifs were involved in regulation of transcription, DNA-dependent and signaling pathway. Based on the results, it seems that the WRKY transcription factor families were the main transcription factors because in the promoter, 85% of the DEGs and the hub genes, involved at least one WRKY binding site. Moreover, these motifs were involved in MFs, including transcription factor activity and protein serine/threonine kinase activity (**Table 4**). GO term analysis of hormonal DEGs showed the motifs involved in regulation of transcription (**Table 5**). The main transcription factor families included WRKY, MYB, BHLH, and bZIP. Finally, this analysis also identified the motifs related to SA pathway (GO: 0009751) (**Table S9**).

### Protein-protein interactions and selection of key genes

It is important to find genes that are regulated for any type of biotic stresses. We selected 1232 biotic DEGs and 308 hormonal DEGs. The DEGs also represent commonly responses to stresses and may be useful in describing general responses to stress in barley and Arabidopsis. For this purpose, the analysis of protein-protein interactions (PPIs) using the STRING database and Cytoscape was performed (**Fig. 9a and b**).

**Fig. 9.**
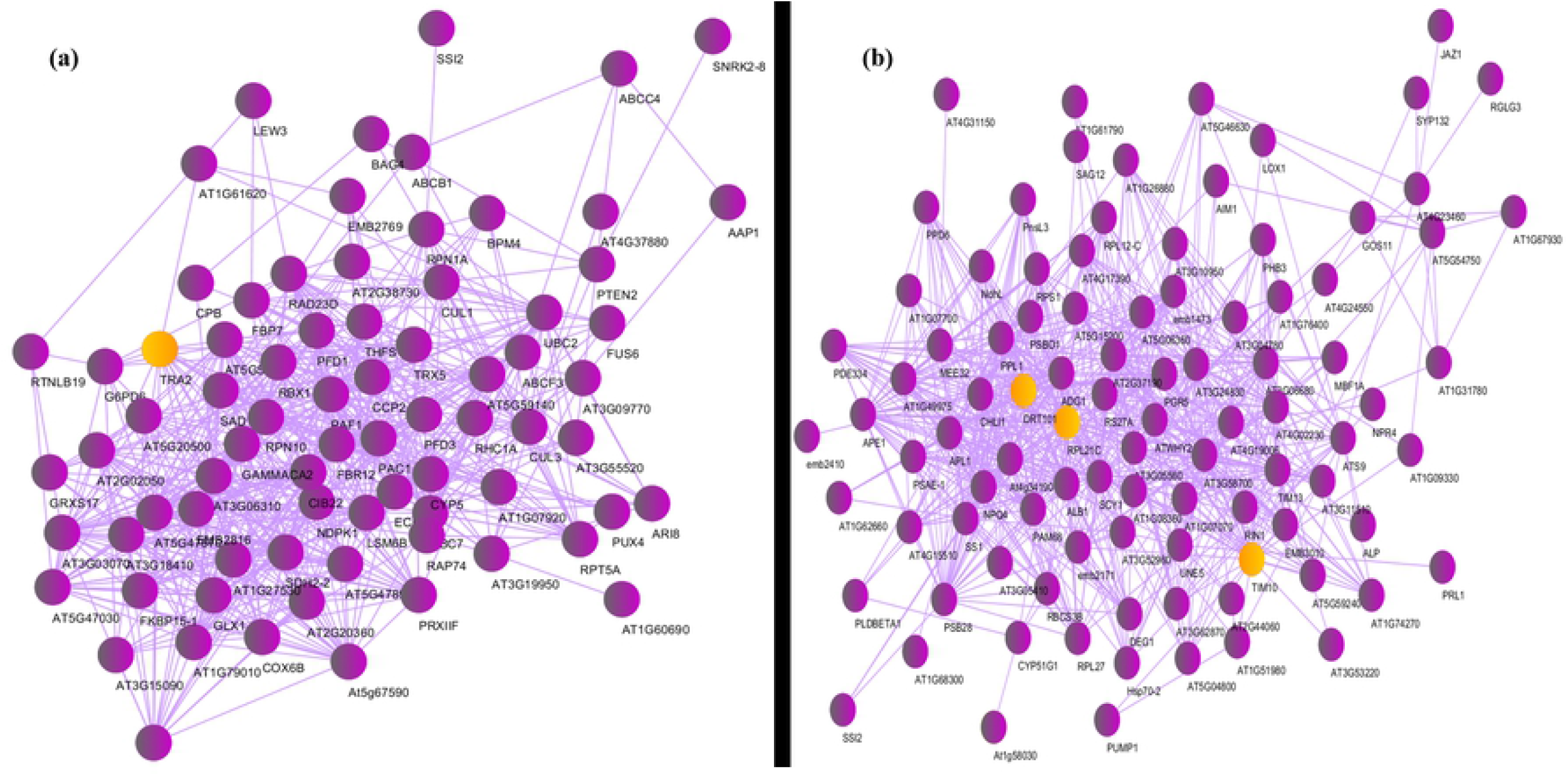
The protein-protein interaction network using STRING database and cytoscape. (a) Network of interactions among biotic hub genes and DEGs. Light colored nodes represent the most connected genes in the network. (b) Network of interactions among hirmonal hub genes and DEGs.

The network was plotted for 265 key biotic DEGs that were involved in the most important mechanisms response to biotic stress and 61 hormonal key DEGs that were related to the hormonal signaling pathways (**Table 6**).

**Table 6.** The most important molecular mechanisms related to biotic and hormonal studies.

| Selective mechanisms related to hormonal data |  | Selective mechanisms related to biotic data |  |
| --- | --- | --- | --- |
| Number of genes | Meaningful mechanism | Number of genes | Meaningful mechanism |
| 15 | Jasmonic acid biosynthetic process | 104 | Oxidation-reduction process |
| 5 | Response to fungus | 9 | Hydrogen peroxide catabolic process |
| 17 | Defense response | 12 | Response to ethylene |
| 3 | Auxin | 6 | Response to reactive oxygen species |
| 6 | Response to karrikin | 11 | Oxidative stress |
| 9 | Defense response to bacterium | 43 | Response to bacterium |
| 2 | Salicylic acid catabolic process | 24 | Response to cytokinin |
| 4 | Response to nematode | 40 | Photosynthesis |
|  |  | 16 | Jasmonic acid biosynthetic process |

A list of genes that showed the most association in the protein interaction networks was then prepared (**Table S10**). Finally, among up-regulated hub genes related to biotic stress and hormonal signaling, it is worthy to notice some of the major genes that may play a key role in fungal response. Therefore, 3 key genes related to biotic stress data including *DNA-DAMAGE-REPAIR/TOLERATION 101* (*DRT101*), active in the DNA repair pathway, *TRANSLOCASE OF THE INNER MEMBRANE 10* (*TIM10*), active in protein import into the mitochondrial inner membrane, and *ADP GLUCOSE PYROPHOSPHORYLASE 1 (ADG1*), active in the starch biosynthetic process, were identified. Among the key hormonal DEGs, the *Aldolase-type TIM barrel family protein* (*TRA2*), active in the response to cadmium ion pathway, was identified. Finally, a PPI network was developed to study key genes that are commonly expressed among biotic and hormonal DEGs. There were a total of 63 common DEGs such as *PAL, NINJA, JAZ1, APE1, RPL24* and some others significant genes especially with unknown function (**Fig. 10**).

**Fig. 10.**
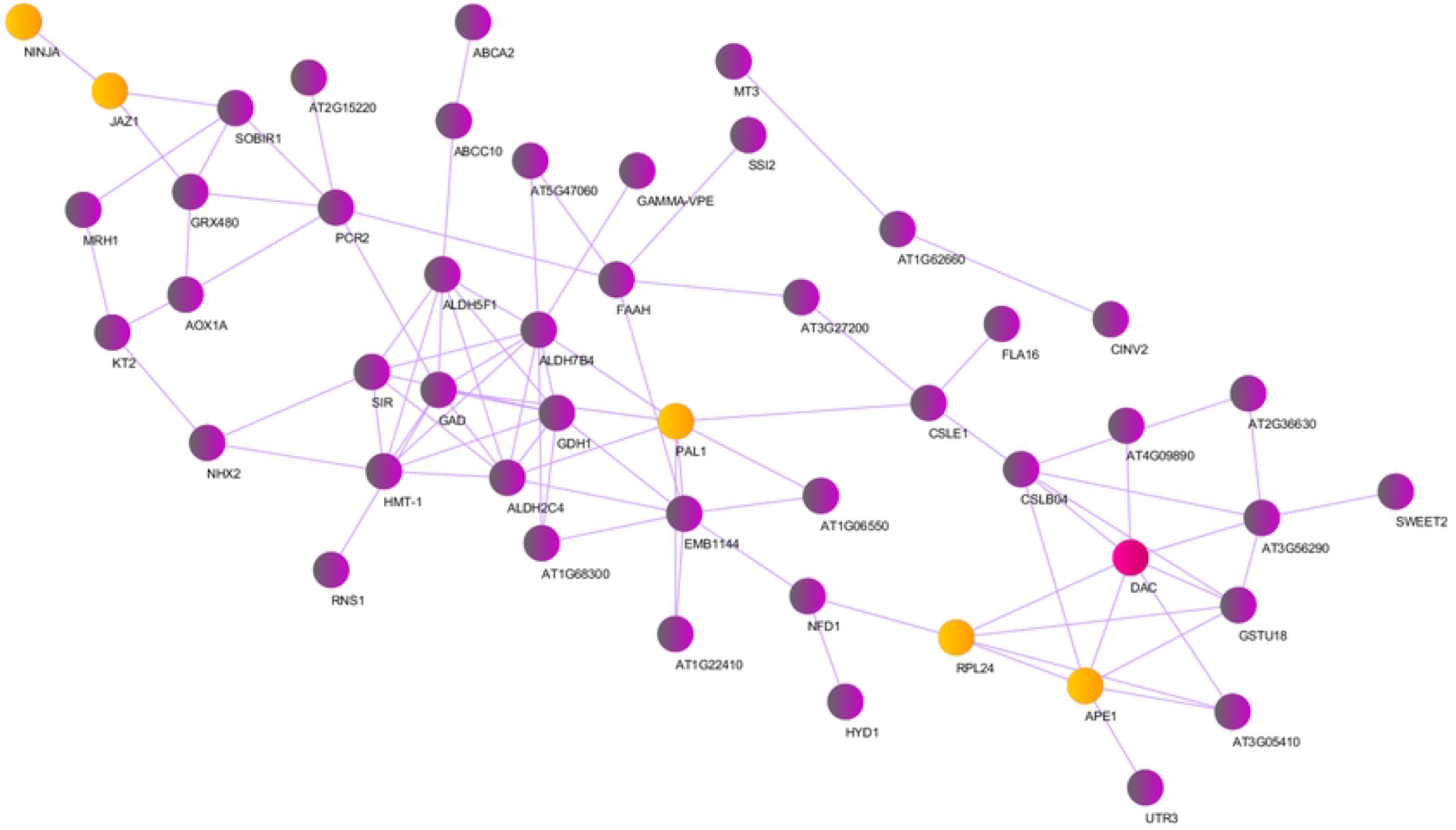
The protein-protein interaction network using STRING database and cytoscape. Interaction network of common genes between hormonal and biotic studies.

## Discussion

Cereals are continuously subjected to biotic stresses, which causes changes in plant metabolism involving physiological damages that leading to a reduction of their productivity. Therefore, understanding plant defense mechanisms prevents significant crop and economic losses (B. Liu *et al*., 2014; Sharma et al., 2013). To fight biotic stresses in barley, the plant has developed complex defense mechanisms which can interact with each other, through the activation of TFs, PKs, microRNAs, and other critical signaling regulators. In addition, the plant defense against pathogens are the result of a series of consecutively changes at the cellular level mainly related to hormone-dependent signal pathways. Resistance induced in plants often appears as an increase in the expression of plant defense genes against stresses and increases plant defense capacity.

In spite of many researches, there has not been sufficient clearance of defense molecular mechanisms for example how are the hormonal signaling cross-talking against various pathogens or what are the main common and specific gene networks. Therefore, one of the most efficient and commonly methods of extracting biological information is the use of transcriptome, which has created a significant ability in researchers to obtain huge amount of data. According to the goal of researchers in using the microarray technique, which actually is to identify different expression genes and function of genes in cell activity, the normalization method that ensures the reliability of this data is also of great importance (Stafford, 2008). In this study, using two different microarray platforms, Affimetrix and Agilent, different within array and between array normalization methods, were compared in order to eliminate or reduce batch effects. It is clear that comparing different process normalization strategies is difficult because their performance depends on a wide range of parameters such as experimental design, data collection process, and image processing software (Y.-J. Chen et al., 2003). Studies by Wu et al. (2005) and Wang et al. (2011) have pointed out that in most transcriptome data with significant genome biological changes, methods such as Quantile do not work well due to hard assumptions (Dong Wang et al., 2011; W. Wu et al., 2005). But our results showed that methods such as Quantile work well for plant transcriptome data normalization (**Fig. S1**).

To identify the DEGs and key pathways, the integration of transcriptomic data received from various biotic stresses and hormonal treatments studies in barley was applied by meta-analysis. Using this approach, a set of components responsible for biotic tolerance and hormone signaling was identified that allow us to understand which genes play a key role in various biotic stresses and hormonal cross-talking. In this way, we collected 479 microarray samples from different biotic stress studies and 46 microarray samples from hormonal treatment studies available in public databases (**Table S1**). In total, the meta-analysis identified 1232 and 304 DEGs for biotic and hormonal studies respectively (**Fig.2 a-b, Table S2**). Of these common DEGs between biotic and hormones studies, 1.2% were categorized as up-regulated and 0.8% as down-regulated (**Fig.3**). These observations showed that under biotic stresses and hormones, a wide range of BPs were up-regulated.

The functional enrichment analysis suggested that DEGs were enriched on various BPs (**Fig. 4-5**). The terms photosynthesis, response to cadmium ion, protein folding, oxidation-reduction process, response to cytokinin, defense response to bacterium, defense response to fungus, cellular response to oxidative stress, response to ABA, JA mediated signaling pathway, response to reactive oxygen species, response to wounding, and response to biotic stresses and hormones were also found to be significant. Carbohydrate metabolism has been mostly shown as an essential pathway affected by biotic stress responses in plants (Martinelli *et al*., 2012). Growing tissues may be seen as a complex of source sinks of carbohydrate attracting photosynthesis produced by leaves. Because the correct mechanism causes the allocation of carbon during stresses consequently to improves plant performance in stress environments (Falchi *et al*., 2020). The sources disruption has been linked with the early pathogenic mechanisms of diseases in plants (Punelli *et al*., 2016). In fact, the dysregulation of this pathway at transcriptomic level may be associated with a general plant stress state. This altered transcript condition may be seen by manufacturers as a sort of “alarm bell” to help further monitoring actions and the initiate of management procedures. In addition, sugar alcohols are directly linked with stress responses. Although their role in tolerance to stress has been more connected with abiotic stresses than biotic ones, it has been hypothesized that they may play an essential role also in a beneficial modulation of biotic stress responses (Balan *et al*., 2018).

It is very essentially to study the plant hormonal responses to pathogens because the signaling pathways of various hormones regulate biotic stress responses antagonistically (Balan et al., 2018). It is observed that several genes (such as WRKY families) involved in auxin, ET, JA, and SA were usually affected in all types of pathogens. Fungal pathogens up-regulate ET, benzyl-adenine (BA), and SA while mostly repress JA-related genes. For example, *Erwinia amylovora* up-regulates ET and GA signaling genes while viruses affect some essential genes involved in ET and auxin signaling (Balan et al., 2018). Our results in this study confirmed that the hormone-related pathways and therefore their crosstalk was deeply affected by all barley pathogens.

To present information regarding how genes regulate in barley against pathogens attack and hormone treatments, we identified TFs as regulatory molecules that play an essential role in gene transcription. Transcription factors not only act as a molecular activator for the expression of stress-inducible genes, but also play a major role in signal transduction pathways in biological networks and promote plant adaptation to various environmental conditions (Tahmasebi *et al*., 2019a). There were 24 TFs for biotic studies belonged to 25 TF families and 6 TFs for hormonal studies belonged to 6 TF families (**Table S5**). The major members of TF families such as GNAT, NF-YC, and Whirly were the top classes and were also considered as hub genes (**Fig. 7**). In this analysis WHIRLY was up-regulated. Plants have a small family of single-stranded DNA-binding proteins called WHIRLY, which are also involved in implicated in defense gene regulation (Desveaux et al., 2004; Vickers, 2017). In the barley, WHIRLY binds to the promoter of the HvS40 senescence-related gene, which is produced during defense against pathogens, during senescence and also hormones such as SA, JA, and ABA, which are synthesized in plastids (Krupinska *et al*., 2014; Lin *et al*., 2020). Transgenic barley with suppressed WHIRLY1 levels remained almost green phenotype despite having higher SA levels than the wild-type (Krupinska et al., 2002). One out of one NF-Y was down-regulated. In plants, NF-Y subunits are encoded by multigene families whereas members show structural and functional diversity. It has been shown that an increasing number of NF-Y genes that play the major roles in the various stages of root node coexistence and arbuscular mycorrhizae as well as in the interaction of plants with pathogenic microorganisms (Hanemian *et al*., 2016). Barley NF-Ys comprise a relatively diverse gene family involved in various processes such as starch metabolism, sucrose metabolism, and photosynthesis. There is less evidence about the role of NF-Y in defending against pathogenic attacks. Overexpression of rice *NF-YA* (*OsHAP2E*) causes resistance to *Magnaporthe oryzae* and *Xanthomonas oryzae* (Alam et al., 2015). In general, the signaling pathways including the *NF-YAs* can regulate beneficially pathogenic plant and microbial interactions and indicate the existence of common mechanisms between both types of biological interactions. These findings corroborate contrast action between regulators that allow plants to create a complex network of interacting pathways in encounter with various stresses and hormonal signaling.

PKs have evolved as the largest family of molecular switches that regulate protein activities related to almost all basic cellular functions. Only a fraction of plant PKs, however, have been functionally characterized even in model plant species. To investigate signal transduction mechanisms in this study, we identified potential PKs that play key roles in signaling networks and plant responses to various biotic stresses and hormones. We detected 3 PKs among the biotic DEGs and 3 PKs among the hormonal DEGs that are classified into RLK-Pelle and TKL groups (**Table 3**). RLK/Pelles play important roles from growth regulation to defense response, and the dramatic expansion of this family has been considered very important for plant-specific adaptations (Ye *et al*., 2017). We realized that hundreds of *RLK/Pelles* are up-regulated by biotic stresses. In addition, the rate of stress responsiveness is related with the degree of consecutive duplication in RLK/Pelle subfamilies (Lehti-Shiu et al., 2009). Members of this family are involved in many different plant processes including regulation of meristem proliferation, reproduction, hormonal signal transduction, and response to biotic and abiotic stresses (Acharya et al., 2007; Florentino et al., 2006; Miya et. al., 2007; Zipfel et al., 2006). Accordingly, these results suggest that PKs play a central role in signal transduction during plant growth and development and also in plant responses to biotic stresses. This suggests that there is a positive and negative cross-talk among PKs.

Plants have evolved complex mechanisms to afford with a diversity understanding small RNA-guided stress regulatory networks that can provide new insights for the genetic improving plant stress tolerance. MicroRNAs (miRNAs) are a major class of small non-coding RNAs that have recently emerged as important regulators of gene expression, during or after transcription (Djami-Tchatchou et al., 2017). Some of miRNAs have been related to biotic stress responses, and the play important roles in plants infected by pathogenic bacteria, viruses, nematodes, and fungi (Anjali & Sabu, 2020). Barley is one of the most important agricultural crops that has been considered for miRNA studies (Djami-Tchatchou *et al*., 2017). In the present study, 432 miRNAs belong to 56 miRNA families for biotic studies and 85 miRNAs belong to 32 miRNA families for hormonal studies and also 15 common miRNAs for both studies were predicted (**Table S6**), a high proportion of which are involved in the different BPs (**Fig. 8b**). The major members of the miRNAs for the both studies belong to conserved families including miR6192, miR6180, and miR6214, miR6196, miR6184, miR166, and miR156. We detected 100 miRNAs for the first time. Many microRNAs (miR156, miR159, etc.) had orthologs in wheat or rice that were recognized to be expressed, while up to a number of microRNAs (miR444) were appeared to be specifically expressed in barley. A combination of small RNA and mRNA degradome analyses to recognition and characterization of many conserved miRNAs (miR156, miR166, etc.) and novel miRNAs (hvu-miR1120b) together with a number of miRNA target genes were done. The results showed that some miRNAs are involved in different functions including carbohydrate translocation, cell differentiation, defense response, photosynthesis, and phytohormone signaling pathways (Shuzuo et al., 2012). This finding revealed that miRNAs play a significant role in regulating early growth and development of the barley and also in the signaling pathways of different hormones that regulate biotic stress responses (Curaba et al., 2012).

Analysis of promoter region of genomic sequences is mainly based on the identification of regions of the genome that are capable of transcription factors binding. The study of these regions and their transcription factors and downstream genes that are regulated by these proteins (target genes) is a fascinating topic in the field of post genomics and can provide new insights into vital BPs (Lemon & Tjian, 2000; Sandelin et al., 2004). We performed a promoter analysis to identify cis-acting elements located in upstream of DEGs according to whether these are under common regulatory control. Eleven with significant scores were identified (**Table S8**). Many cis-regulatory elements are related with cross-talking to pathogens and hormones. Many motifs, which was found out of DEG promoters, are highly matched to the MA1267.1 motif (**Table S7, Tables 4 and 5**) that is one of cis-acting elements of C2H2 zinc finger factors (DOF5.8). DOF TFs are zinc-finger regulators distributed among non-vascular and vascular plants. DOF proteins play an important role in many BPs as, e.g., flowering time, seed development, and responses to hormones and stresses (Ruta *et al*., 2020). In addition, He et al. realized the Arabidopsis DOF5.8 to be an upstream regulator of a gene encoding an NAC family member in response to stress (Klees *et al*., 2021). More particular, DOF5.8 was shown to play a major role in the correct development of veins in the leaves of *Arabidopsis thaliana* and its promoter was demonstrated to be regulated by auxin (Konishi *et al*., 2015). Actually, the overexpression of this TF leading to a modification in the expression of many genes was involved in auxin biosynthesis, vascular tissue formation, secondary cell wall deposition and as well as signaling during physiological processes (Guerriero *et al*., 2020; Konishi & Yanagisawa, 2015).

Then, co-expression network analysis of the DEGs was performed to discover processes and functions that are involved in response to pathogens and hormonal signaling. We recognized 6 significant modules in biotic studies and 7 significant modules in hormonal studies as principal functional clusters (**Fig. S2**). Enrichment analysis of modules in both studies showed a wide range of BPs related with signaling, photosynthesis, defense response to bacterium or fungus and response to JA (**Table S3**). Interestingly, for both studies, 4 modules including Brown, Blue, Turquoise, and Yellow were significantly related with response to stresses. After studying of the genes within these modules, finally three genes (*ADG1, TIM10*, and *DRT101*) related to biotic studies and one gene (*TRA2*) related to hormonal studies that were involved in the regulation of BPs to further investigation were identified (**Table S10, Fig. 9**).

Starch biosynthesis plays a central role in plant metabolism, so starch is an important determinant of the growth and development of non-aerial organs (Kishore, 1994). Several enzymatic stages are involved in starch biosynthesis in plants. ADP-glucose pyrophosphorylase (ATP: alpha-glucose-1-phosphate adenylyl transferase, ADGase) has been studied as a major regulatory enzyme in the starch biosynthetic pathway in plants (Crevillén et al., 2005; S. M. Wang et al., 1998b; Zeeman et al., 2007). There are 6 genes encoding proteins with homology to ADP-GlcPPase, 2 of these genes encode small subunits (APS1: ADG1 and APS2). Surprisingly, *ADG1* can be expressed and identified not only in chloroplast but also in non-plastidic regions, especially small proportion in nucleus (Bahaji et al., 2011). Genetic elements and molecular mechanisms leading the flowering process in *Arabidopsis thaliana* flowering plants and short-day flowering rice have been investigated (Cho et al., 2017). Mutants of *Arabidopsis thaliana* starch metabolism with *ADG1* (AT5G48300) showed higher sugar concentrations the leaves and exhibit late flowering phenotypes (Caspar et al., 1991; Eimert et al., 1995; Lin et al., 1988). In addition, penetration resistance indicates the first level of plant defense against phytopathogenic fungi. Therefore, it has been reported that the starch-deficient *Arabidopsis thaliana ADG1* mutant has impaired penetration resistance against the hemibiotrophic fungus *Colletotrichum higginsianum* (Engelsdorf et al., 2017). The protein interaction network for the *ADG1* was drawn using STRING software in which 11 genes with 54 interactions participated in this network (**Fig S5a**). The results showed that 9 genes involved the highest interaction with other genes. One of the most important genes in this network is the *STARCH SYNTHASE 1 (SS1*), which, if presents in the plant is broken down the Cys bridge between *ADG1* and activates the ADP-glucose pyrophosphorylase, which is responsible for starch synthesis. Plants have evolved a set of antioxidant defense systems to defend themselves against a host of stress reactions (Yang et al., 2018). The *DRT101* (DNA-damage-repair/tolerance protein) encoding important enzymes responsible for antioxidant defense was identified. This gene was strongly expressed in auxin synthesis, cell division, and antioxidant defense (Bian et al., 2019). Studies showed that DRT proteins (Fujimori et al. 2014) are assigned with defensive functions for scavenging the UV-B-induced DNA damage against sunlight and pathogens. The studies have been reported that the DRT101 is nuclear-encoded but chloroplast-targeted (Bian *et al*., 2019). The protein interaction network for *DRT101* in which 11 genes with 33 interactions participated was showed (**Fig. 9b**). One of the most important genes in this network is the transcription factor of GNAT that we identified in the both biotic stress and hormonal pathways. TIM10 is a small protein from the TIM chaperone complex (**Fig. S5c**) that facilitates the transfer of hydrophobic proteins into the inner mitochondrial membrane and aids to protein folding. The assembly of TIM10 also depends on oxidative stress in mitochondria and messaging molecules. TIM10 is loaded with Zn^2+^ in the cytosol. The role of Zn^2+^ is to keep the protein in the reduced and unfolded form in the cytosol, before entering into the mitochondria (Herrmann & Neupert, 2000). In Arabidopsis, there are two genes encoding TRA (*TRA2*, *TRA1*). TRA2 (**Fig. S5d**) that is involved to maintain ROS balance in response to Glc during early seedling growth and sheds light on the relationship between Glc, the pentose phosphate pathway and ROS. In general, TRA activity is related to plant defense mechanisms, which include the synthesis of secondary metabolites and is one of the fastest responses to pathogenic attacks in plants, as well as to programmed cell death and wounding of injured tissues (Zheng *et al*., 2020).

## Conclusion

In this study, we integrated a meta-analysis method with the gene co-expression network analysis on microarray studies to identify the DEGs involved in biotic stresses, hormonal treatments, and some cross-talking in barley. This type of approach increases sensitivity in the identification of important stress response genes that may be missed by studies that are limited to specific tissue or growth stage or level of stress. Most of the DEGs were involved in pathways of photosynthesis, response to cadmium ion, protein folding, oxidation-reduction process, defense response to bacterium, defense response to fungus, cellular response to oxidative stress, JA mediated signaling pathway, and response to reactive oxygen species. In addition, using DEGs, various analyzes including analysis of promoter, identification of PKs and transcription factors were performed. These analyses extend the role of TFs (e.g., WRKY, MYB, BHLH, and bZIP), kinases (RLK/Pelle), miRNAs (e.g., miR6192) which are key to the pathogen response, while a number of novel genes with unknown functions were also identified. Results help us for more deep insight on molecular mechanisms against different pathogens and on hormonal cross-talks. Weighted gene co-expression network analysis divided genes into individual and consensus modules and showed sets of genes with conserved and reversed expression status. A number of genes with high connectivity and conserved expression but with weak annotation were recognized. We suggest these DEGs, which might provide a more robust bio-signature for phenotypic traits, as a more promising resource of molecular biomarkers and potential candidate genes for engineering pathogen tolerance in barley.

## Acknowledgments

The authors would like to thank the Institute of Biotechnology for supporting this research and the Bioinformatics Research Group in the College of Agriculture (Shiraz University).

## Declarations

The authors declare that there is no conflict of interest.

